# Develop an efficient and specific AAV based labeling system for Muller glia in mice

**DOI:** 10.1101/2021.12.10.472182

**Authors:** Yanxia Gao, Kailun Fang, Zixiang Yan, Haiwei Zhang, Guannan Geng, Weiwei Wu, Ding Xu, Heng Zhang, Na Zhong, Qifang Wang, Minqing Cai, Erwei Zuo, Hui Yang

## Abstract

Cell degeneration in the retina leads to several ocular diseases and vision loss. Considerable research efforts focus on reprogramming Muller glia (MG) into functional cells to rescue vision as a promising therapeutic strategy, although whether MG can convert into functional bipolar cells, retinal ganglia cells (RGCs), rods or cones in mammals remains controversial. The broad applicability of tracking MG differentiation thus presents a need for improved labeling efficiency and specificity. Here, we compared AAV-based labeling strategies with conventional lineage-tracking by crossing transgenic mouse lines. We found that reporter expression was weak and not MG-specific in *mGFAP-Cre* transgenic mice. Different AAV serotypes showed a range of efficiency and specificity in labeling MG, leading us to optimize *a human GFAP-Cre* reporter system packaged in the AAV9 serotype with the *WPRE* (WPRE, woodchuck hepatitis virus post-transcriptional regulatory element) removed. The *hGFAP-Cre-* Δ*WPRE* reporter could label 20-73.8% MGs, with non-specific RGC labeling rates ranging from 0-0.08% at doses of 1 × 10^8^ to 10^10^ vector genomes (vg) per eye, an approximate 40-fold reduction compared with the *AAV9-hGFAP-Cre-WPRE* labeling system. Furthermore, we validated the availability of the label system to trace MG reprogramming and also detected false-positive results reported previously. The *AAV9-hGFAP-Cre-*Δ*WPRE* system thus represents a highly efficient and specific labeling system for MG, providing a valuable tool for tracking cell fate *in vivo*.

## Introduction

Muller glia (MG) represent the main type of glial cell which are responsible for maintaining retinal structure and support for neurons in the retina. In lower vertebrates (e.g., zebrafish, *Danio rerio*), MG can re-enter the cell cycle to proliferate and subsequently differentiate into one of multiple cell types following injury, such as photoreceptors or retinal ganglia cells (RGCs) (Bernardos, Barthel, Meyers, & Raymond, 2007; Fimbel, Montgomery, Burket, & Hyde, 2007; Goldman, 2014; Sherpa et al., 2008). However, the process does not occur in mammals. As a result, many ocular diseases that are difficult to treat, such as glaucoma or retinitis pigmentosa, develop from functional degeneration or loss of neurons. Thus, decades of research efforts have been committed to investigating the regenerative machinery in adult mammals with the ultimate goal of inducing MG regeneration and reprogramming. Several major insights have emerged from this work. For example, in mice Ascl1 overexpression or simultaneous deletion of three nuclear factors I (NFI) genes can both induce MG proliferation and transformation into amacrines or bipolar cells (Hoang et al., 2020; Ueki et al., 2015). Similarly, MG to conversion to RGCs was also observed in mice following deletion of Ptbp1 or simultaneous ectopic expression of Pou4f2 (Brn3b) and Atoh7 (Math5) (Xiao et al., 2021; Zhou et al., 2020). Moreover, photoreceptors could be generated from MG under simultaneous overexpression of Otx2, Crx and Nrl after a period of β-catenin ectopic expression (Yao et al., 2018).

However, *in vivo* glial lineage-tracing methods represent a long-standing research objective. For instance, an AAV-mediated glial reporter in the brain has been used by different research groups to observe the conversion of neuroD1-induced astrocytes into neurons (Boutin et al., 2010; Brulet et al., 2017; Chen et al., 2020; Wang et al., 2021). However, this phenomenon was not detectable using a more stringent, inducible, transgenic lineage-tracing strategy (Wang et al., 2021). One proposed explanation is that AAV-mediated reporters also non-specifically label neurons, suggesting that no conversion was observed, or alternatively, that inducible transgene methods cannot label all astrocytes, and thus do not label the resulting reprogrammed neurons. Similar confounding factors have been reported in MG lineage-tracing systems, such as the low or extremely variable labeling efficiency in inducible CreER mouse lines (Jorstad et al., 2017; Rueda et al., 2019; Thanh Hoang et al., 2021; Ueki et al., 2015), or potential non-specific labeling of other cell types by AAV-based methods, similar to the problems in astrocyte studies mentioned above (Yao et al., 2016; Yao et al., 2018). It is therefore crucial to develop an efficient and stringent tool for tracing MG proliferation and reprogramming in the retina.

Here, we compared the labeling efficiency and specificity of two different reporter systems: those based on crossing *mGFAP-Cre* transgenic mice and those based on transfection with AAV-*hGFAP-Cre* constructs in Ai9 mice, which harbor the *Rosa26-CAG-loxp-stop-loxp-tdTomato* inducible reporter. By optimizing the AAV reporters through WPRE (WPRE woodchuck hepatitis virus post-transcriptional regulatory element) cassette deletion, selecting the most suitable serotype, conducting immunofluorescence microscopy to quantify construct specificity and efficiency *in vivo*, we provide a novel, highly specific and reproducible labeling system for MG cells in the retina tissue of mice. By re-examining MG reprogramming with this labeling system through overexpression or downregulation of transcription factors reported previously, we validated its availability and detected some false-positive results.

## Materials and methods

### Animals

All mice were housed under a 12 hour light/dark cycle with water and food provided *ad libitum* at the Animal Center of the Institute of Neuroscience, Chinese Academy of Sciences, Shanghai, China. All animal procedures were approved by the Animal Care and Use Committee. All reporters were based on tdTomato expression in *Ai9* transgenic mice (*CAG-Loxp-stop-Loxp-tdTomato*, JAX #007909) purchased from the Jackson Laboratory. *GFAP-Cre* (B6.Cg-Tg(Gfap-cre)73.12Mvs/J, JAX#012886) mice were gifted by Dr. Jiawei Zhou and *Aldh1l1-Cre* (Tg(Aldh1l1-cre)PB1Gsat/Mmucd) were gifted by Dr. Zilong Qiu in the Institute of Neuroscience, Chinese Academy of Sciences.

### AAVs and vectors

All AAV vectors were constructed by PCR-based subcloning. The 681-bp human GFAP promoter was used in this study, which was derived from the 2.2-kb gfa2 promoter (Y. Lee, Messing, Su, & Brenner, 2008). Different serotypes of AAVs were packaged and titered by Gene Editing Core Facility in the Institute of Neuroscience.

### Subretinal injection

AAVs were delivered to eyes via subretinal injection, as previously described (Chiu, Chang, & So, 2007; Qi et al., 2015). For subretinal injection, adult mice were anaesthetized with a mixture of ketamine and xylazine, and pupils were dilated with tropicamide phenylephrine eye drops before injection. A small scleral incision was made using a 30G needle under a microscope (Olympus, Tokyo, Japan). Then a 32G needle on a Hamilton syringe was inserted into the subretinal space though the scleral incision. 1 μl of AAV was slowly (i.e., up to 20s) injected into the subretinal space. After AAV injection, the Hamilton syringe was removed and a drop of ofloxacin eye ointment was applied to cover the eye. In gene transfer of transcription factors experiments, mice were injected at 4∼8 weeks old.

### Immunofluorescent staining and Imaging

Mouse eyes were collected after perfusing animals with 4% paraformaldehyde (PFA) in PBS, and post-fixed for half an hour in PFA at room temperature after removing the cornea. Then, retinas were dissected and dehydrated overnight using 30% (*w/v*) sucrose in PBS and embedded in O.C.T (Sakura). Sections were performed by a cryostat (HM525, Thermo). Before staining, sections were dried for 30 minutes at 65°C and then were washed in PBS 3 times for 5 minutes per wash. Antigen recovery was performed using heated sodium citrate. Blocking was performed in 150 μl blocking buffer (0.1 M PBS: 10% goat serum, and 0.1% Tween-20). Primary antibodies were incubated at 4°C overnight. Details of primary antibodies used in this study were as follows: rabbit anti-Rbpms (1:200, Proteintech, 15187-1-AP), mouse anti-Brn3a (1:100, Millipore, MAB1585), mouse anti-Sox9 (1:200, Sigma, Amab90795), rabbit anti-Rfp (1:200, Rockland, 600-401-379), Mouse-anti-PKC α (1:200, Santa Cruz Biotechnology, sc-8393), Rabbit-anti-Calbindin (1:200, Proteintech, 14479-1-AP) and mouse-anti-Calretinin (1:200, Proteintech, 66496-L-IG), rabbit anti-Pax6 (1:200, Proteintech, 12323-1-AP). Sections were washed 3x for 10 minutes per wash on the second day. Secondary antibodies were also incubated at 4°C overnight. These secondary antibodies included: Alexa Fluor 647-AffiniPure goat anti-mouse IgG (H+L) (1:500, Jackson ImmunoResearch, 115-605-003), Alexa Fluor 647-AffiniPure goat anti-rabbit IgG (H+L) (1:500, Jackson ImmunoResearch, 111-605-003), Fluorescein (FITC) AffiniPure goat anti-mouse IgG (H+L) (1:500, Jackson ImmunoResearch, 115-095-003), Cy™3 AffiniPure goat anti-rabbit IgG (H+L) (1:500, Jackson ImmunoResearch, 111-165-003). Sections were washed 3x for 10 minutes per wash on the third day. After nucleic DNA staining by 4′,6-diamidino-2-phenylindole (DAPI), sections were mounted with fluorescent anti-fade mounting medium (Southern Biotech, 0100-01). All sections were imaged by confocal microscopy (Olympus, FV3000). Images were processed using ImageJ (Zhou et al., 2020).

## Result

### MGs are not accurately labeled in some strains of transgenic mice

We first sought to test the specificity of MG labeling by *mGFAP-Cre* transgenic mouse, as reported in previous work to label astrocytes (Garcia, Doan, Imura, Bush, & Sofroniew, 2004). To this end, we generated an *Ai9;mGFAP-Cre* mouse line to label GFAP positive cells (figure 1A) and found that these tdTomato-labeled cells were distributed throughout the inner nuclear layer (INL) (Figure 1B, C), whereas MGs were distributed within a relatively tight layer within the middle of the INL (Figure 1B). Immunofluorescence staining showed the tdTomato reporter poorly labeled MGs, since only 6.42% ± 0.59% of total tdTomato^+^ cells (501/7722) were co-labeled with the MG marker Sox9 (Figure 1D). However, we also found that a small proportion of tdTomato^+^ cells scattered in the ganglia cell layer (GCL) (Figure 1B), and these cells did not strictly co-localize with the RGC markers Rbpms or Brn3a (i.e., only overlapped with each marker in five cells, or <2% of total RGCs, per eye section) (Figure 1E-F). These results indicated that the endogenous GFAP-driven reporter did not leak to RGCs. Cumulatively, these results demonstrated that MGs were not accurately labeled by endogenous m*GFAP-Cre* in this mouse line.

**Figure 1.**
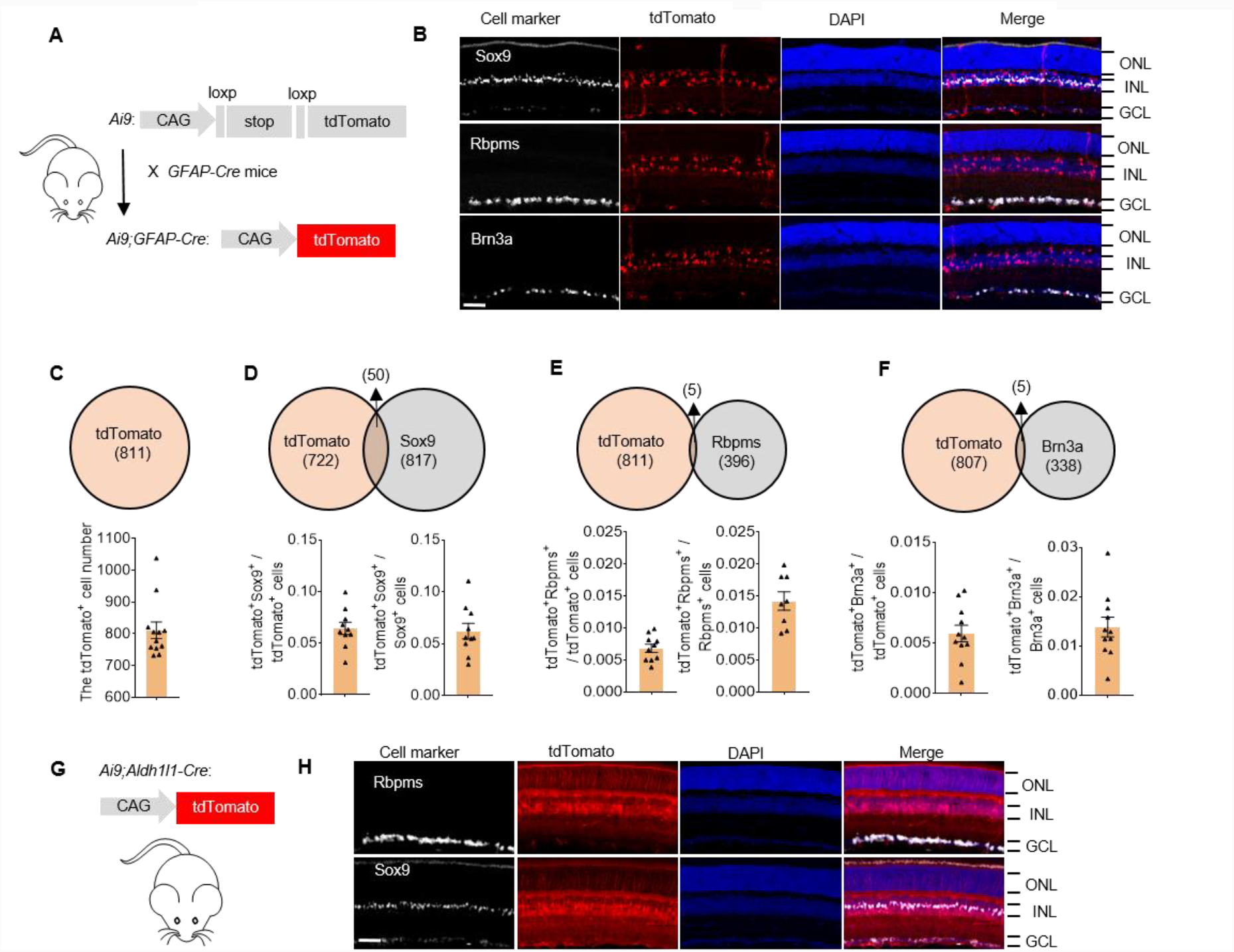
Muller glia are not well-labeled in the *transgenic* mouse line. (A) Schematic illustration of constructs used to establish the *Ai9;mGFAP-Cre* mouse line. (B) tdTomato reporter co-localization with cell markers in *Ai9;mGFAP-Cre* mice. Rbpms and Brn3a are RGC markers; Sox9 specifically labels Muller glia (MG) in the retina. The tdTomato reporter is weakly co-expressed with these cell markers, which indicates poor labeling of MG by tdTomato and no detection of leakage to RGCs. Scale Bar, 50 μm. RGC, retinal ganglia cell; ONL, outer nuclear layer; INL, inner nuclear layer; GCL, ganglia cell layer. (C) The tdTomato cell number in each eye section. n = 4 retinas per group. (D-F) Venn diagram of average number of tdTomato-labeled cells and/or other cell markers in each section. n = 4 retinas per group. All values are presented as means ± S.E.M. (G) Schematic illustration of the *Ai9;Aldh1l1-Cre* mouse line. (H) tdTomato reporter co-localization with cell markers in *Ai9;Aldh1l1-Cre* mice. Scale Bar, 50 μm.

Besides, we also generated an *Ai9;Aldh1l1-Cre* mouse line to label GFAP positive cells (figure 1G) and found that a proportion of tdTomato-labeled cells were not co-labeled with Sox9, although labeled cells did not co-localize with Rbpms (Figure 1H). These results indicated that the endogenous Aldh1l1-driven reporter also labels other types of cells in INL, besides MG cells. In all, these two transgenic mouse lines were not proper to label MG cells.

### AAV-based reporter labels MGs efficiently but not specifically

AAV-based Cre expression is also commonly used to label MG cells in mice due to its relatively easier introduction compared to generating transgenic mouse lines (Xiao et al., 2021; Yao et al., 2016; Yao et al., 2018; Zhou et al., 2020). However, the efficiency of AAV-based labeling also depends on transfection efficiency, which can be affected by AAV serotype. To evaluate the labeling efficiency of different AAV serotypes, we packaged the *hGFAP-Cre-WPRE* construct into commonly used AAV vectors, including AAV1, AAV2, AAV5, AAV8, AAV9, and AAV.ShH10 (Buck & Wijnholds, 2020; Yao et al., 2018), and separately delivered them to the retinas of Ai9 mice by subretinal injection (Figure 2A). The results showed distinct patterns of transfection for the different serotypes. In particular, AAV1 and AAV5 showed less diffusion in the retina and their transfection area was smaller than that of other vectors, while AAV8 and AAV9 exhibited higher transfection efficiency than other vectors, with >500 labeled cells per section, on average (Figure 2B-D). Unexpectedly, the tdTomato reporter showed highly variable leakage (i.e., RGC labeling) among serotypes (Figure 2E). Notably, AAV2 and AAV.ShH10 labeled a greater number of RGCs (>50, or 10% of total tdTomato^+^ cells on average per eye section) than other vectors (Figure 2F-G). These results implied that the selection of specific AAV serotypes could improve transfection efficiency, but could not prevent non-specific labeling of RGCs (or possibly other cell types). Thus, an alternative method was necessary to improve labeling specificity.

**Figure 2.**
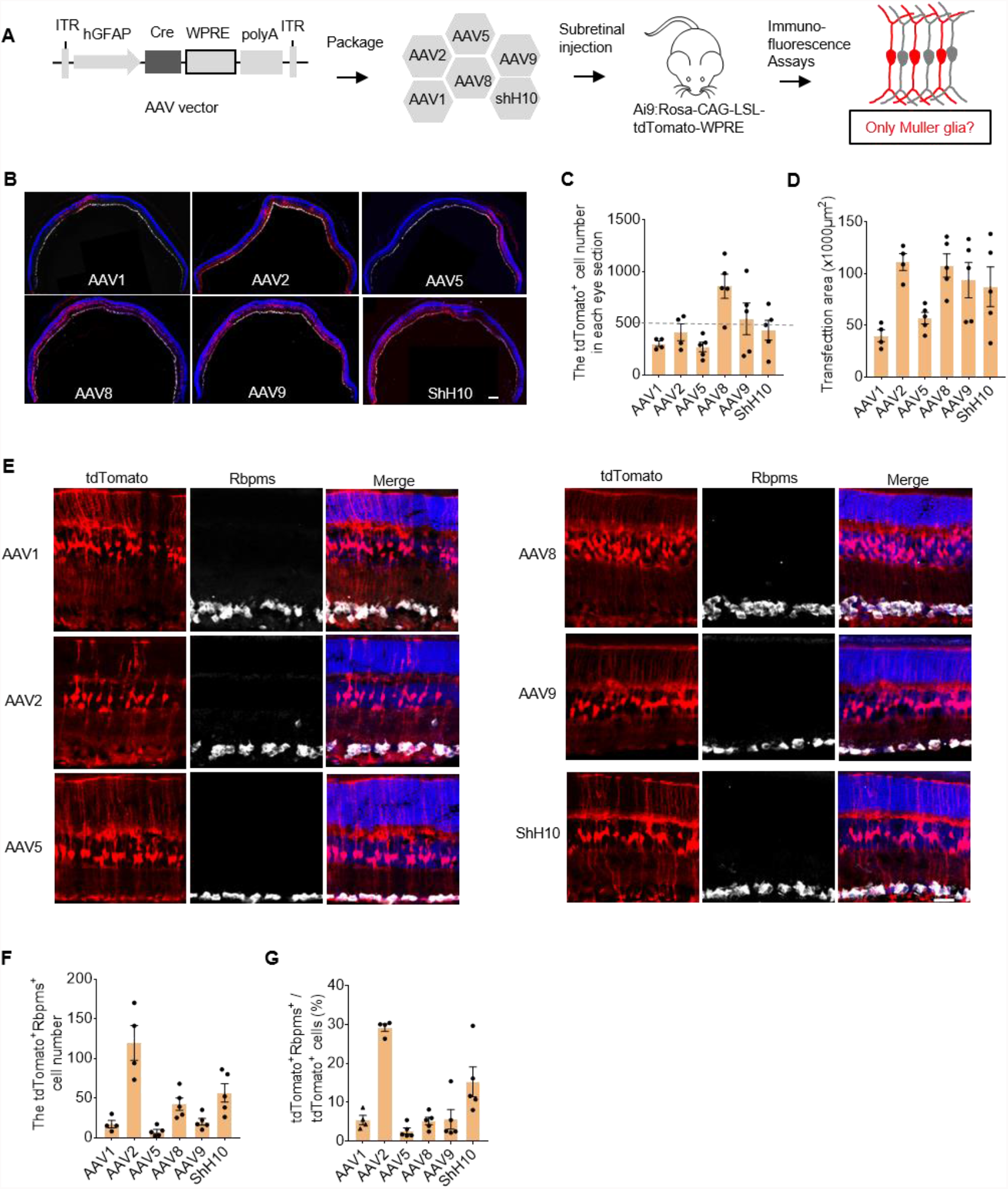
AAV-based systems co-label a small proportion of RGCs along with MG. (A) Schematic illustration of experimental design. Cre expression is driven by the human GFAP promoter and enhanced by the *WPRE* element. WPRE, woodchuck hepatitis virus post-transcriptional regulatory element. (B) Representative images of transfection with different AAV serotypes through subretinal injection. Scale bar, 200 μm. (C) The average number of tdTomato-labeled cells in each eye section infected by different AAVs. (D) Quantification of transfection area in each section. (E) Representative images of co-localization of tdTomato signal with Rbpms in the retina for each AAV serotype. Scale Bar, 50 μm. (F) The number of tdTomato-labeled RGCs. (G) The ratio of tdTomato-labeled RGCs to total labeled cells. n = 4∼5 retinas per group for panels C, D, F and G. All values are presented as means ± S.E.M.

### Deletion of the *WPRE* element in the AAV9 construct eliminates non-specific neuron labeling

Since *WPRE* is often used to enhance gene expression in AAV vectors (Challis et al., 2019; Choi et al., 2014; Deverman et al., 2016), we hypothesized that increasing Cre expression could lead to tdTomato activation in cells with low GFAP promoter activity, potentially resulting in non-specific labeling of RGCs by AAV-based reporters targeting MG. To test this hypothesis, we deleted *WPRE* from the reporter construct (Figure 3A) and packaged it in the AAV9 serotype for delivery by subretinal injection to the eyes of Ai9 mice. We then evaluated the efficiency and MG specificity of tdTomato+ labeling. We found that transfection area and cell number varied in a dose dependent manner, with more extensive reporter signal in the high dose injection groups (1×10^9^ and 1×10^10^ vector genomes /eye) and smaller transfection area and fewer labeled cells in the 1×10^8^ group (Figure 3B-C). However, transfection area and cell number were comparable between the *WPRE* and Δ*WPRE* groups at 1×10^9^ dose (Figure 3C). Among total tdTomato^+^ cells, an average of 700 (73.8% ± 8.5%), 423 (51.6% ± 9.4%), and 153 (20% ± 4.16%) cells were co-labeled with the Sox9 marker in the 1×10^10^, 1×10^9^, and 1×10^8^ groups, respectively, accounting for 99%, 98%, and 96% of the total labeled cells seprately (Figure3 E-F). These results indicated that deletion of *WPRE* could keep the efficiency and improve the selectivity of in vivo MG labeling.

**Figure 3.**
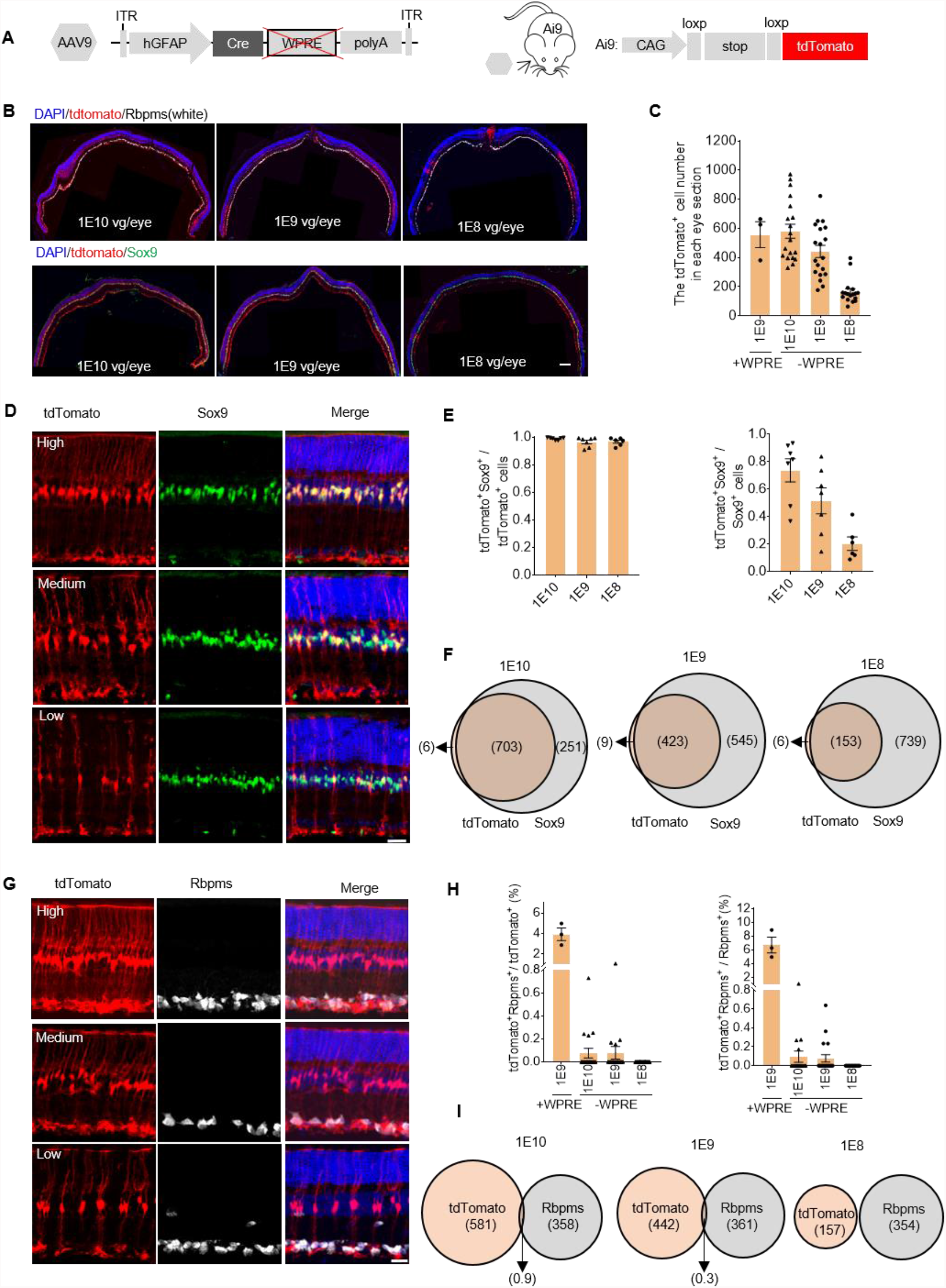
Reduced non-specific labeling in the *ΔWPRE* system. (A) Schematic diagram of the *hGFAP-Cre-*Δ*WPRE* vector and the tdTomato reporter construct in the Ai9 mouse line. (B) Representative images of tdTomato co-localization with the Muller glia marker Sox9 in mouse retinas following subretinal injection with different doses of AAVs (from left to right:1×10^10^, 1×10^9^, 1×10^8^ vector genomes per eye, vg/eye). Scale bar, 200 μm. (C) The average number of tdTomato^+^ cells in each eye section infected by different doses of AAVs. (D) Representative images of tdTomato co-labeling with the Sox9 MG marker in different densities of labeled MG in retina sections. Scale Bar, 50 μm. (E) The ratio of tdTomato-labeled MG among total labeled cells (left) and MG (right). (F) Venn diagram of the average number tdTomato-labeled cells and MG in each section. (G) Representative images of MG cells in retina sections co-labeled with tdTomato and RGC marker Rbpms in different transfection densities. Scale Bar, 50 μm. (H) The ratio of tdTomato-labeled RGCs among total reporter cells (left) and RGCs (right). (I) The Venn diagram of average tdTomato-labeled cells and RGCs in each section. n = 3∼7 retinas per group for panel C, E, H. All values are presented as means ± S.E.M.

Moreover, tdTomato^+^Rbpms^+^ cells were largely undetectable in each of the different AAV dosage groups, even in high density tdTomato^+^ cells (Figure 3G), and with strong statistical significance, since less than one RGC (15 tdTomato^+^Rbpms^+^ cells in 22269 total labeled cells) on average was labeled per eye section (Figure 3H-I). Under the same dose (1×10^9^ vg/eye), the Δ*WPRE* group showed a 40-fold reduction in the ratio of tdTomato^+^Rbpms^+^ cells to total tdTomato^+^ cells (from 3.93% ± 0.61% to 0.08% ± 0.05%), and an 80-fold (from 6.72% ± 1.15% to 0.08% ± 0.04%) among total Rbpms^+^ cells (Figure 3H), compared with the *WPRE* group. It warrants mention that the proportion of labeled RGCs remained consistently low across a wide dosage range, 0.08% to 0% from 1×10^10^ to 1×10^8^ vg per eye in the Δ*WPRE* group. Collectively, these results indicated that the *ΔWPRE* AAV reporter system could labeled MG with high specificity, with only negligible staining of RGCs, even in experiments requiring high titer transfection.

We observed that a very small proportion of tdTomato^+^ cells were not labeled by Sox9 marker for MG (e.g., 9/431 in the 1×10^9^ group in one section, Figure 3G), leading us to question whether these cells were other neuron types, such as bipolar cells, amacrine cells or horizontal cells. To explore this possibility, we conducted immunofluorescent staining for PKC-α (the bipolar cell marker), Calbindin (marked horizontal cells, amacrine cells and RGCs, but could distinguish horizontal cells in mice) and Calretinin (mainly marked mouse amacrine cells) (Kovacs-Oller et al., 2019; Liets, Eliasieh, van der List, & Chalupat, 2006) in eye sections with tdTomato^+^ cells labeled by the Δ*WPRE*-AAV9 system and detected no co-labeling among these cells. (Figure S1A-D). However, we found a very small proportion of tdTomato^+^ cells (<11.8%) were co-localized with Pax6, another amacrine marker (Figure S1E). These results indicated that this reporter did not label other retina neuron types, except for amacrines, either. Collectively, these findings demonstrated that a *AAV9-hGFAP-Cre-ΔWPRE* reporter system could be used to label MG with high specificity and at a wide dosage range in Ai9 mouse retina.

### The Δ*WPRE* system can trace MG reprogramming

To validate *AAV9-hGFAP-Cre-ΔWPRE* reporter system, we traced *in vivo* MG conversion to RGC with the Δ*WPRE* system and *hGFAP-GFP*. Combining with label systems, we performed gene transfer of transcription factors math5 combining with Brn3b, and NeuroD1, reported in previous work to induce MG conversion to neuron and reprogramming (Figure 4A). We found most of tdTomato-labeled cells overexpressed transcription factors, more than 73.8% at 2 weeks after injection and more than 94.4% at 4 weeks (Figure 4B-D). However, no tdTomato^+^Rbpms^+^ cells were detected in both overexpression and control groups at 2 weeks after injection, and the ratio of tdTomato^+^Rbpms^+^ to tdTomato^+^ cells in overexpression groups was low as the control group at 4 weeks (Ctrl:0/2435, Math5/Brn3b: 2/2270, NeuroD1: 3/1839) (Figure 4B, C, E). But we detected several GFP+Rbpms+ cells (126/1843 at 2 weeks, 135/1993 at 4 weeks) in the Math5/Brn3b group, much less in the control and NeuroD1 groups (Figure 4B, C, F), as previous work reported (Xiao et al., 2021). These results suggested the *hGFAP-GFP* label system may label resident RGC mediated by Math5/Brn3b. Besides, we knocked Ptbp1 down by Cas13X.1(Xu et al., 2021) in MG but also did not detect tdTomato^+^Rbpms^+^ cells in the GCL (Figure 4 F-H). These results suggested conversion of MG to RGC fails to be traced mediated by these transcription factors.

**Figure 4.**
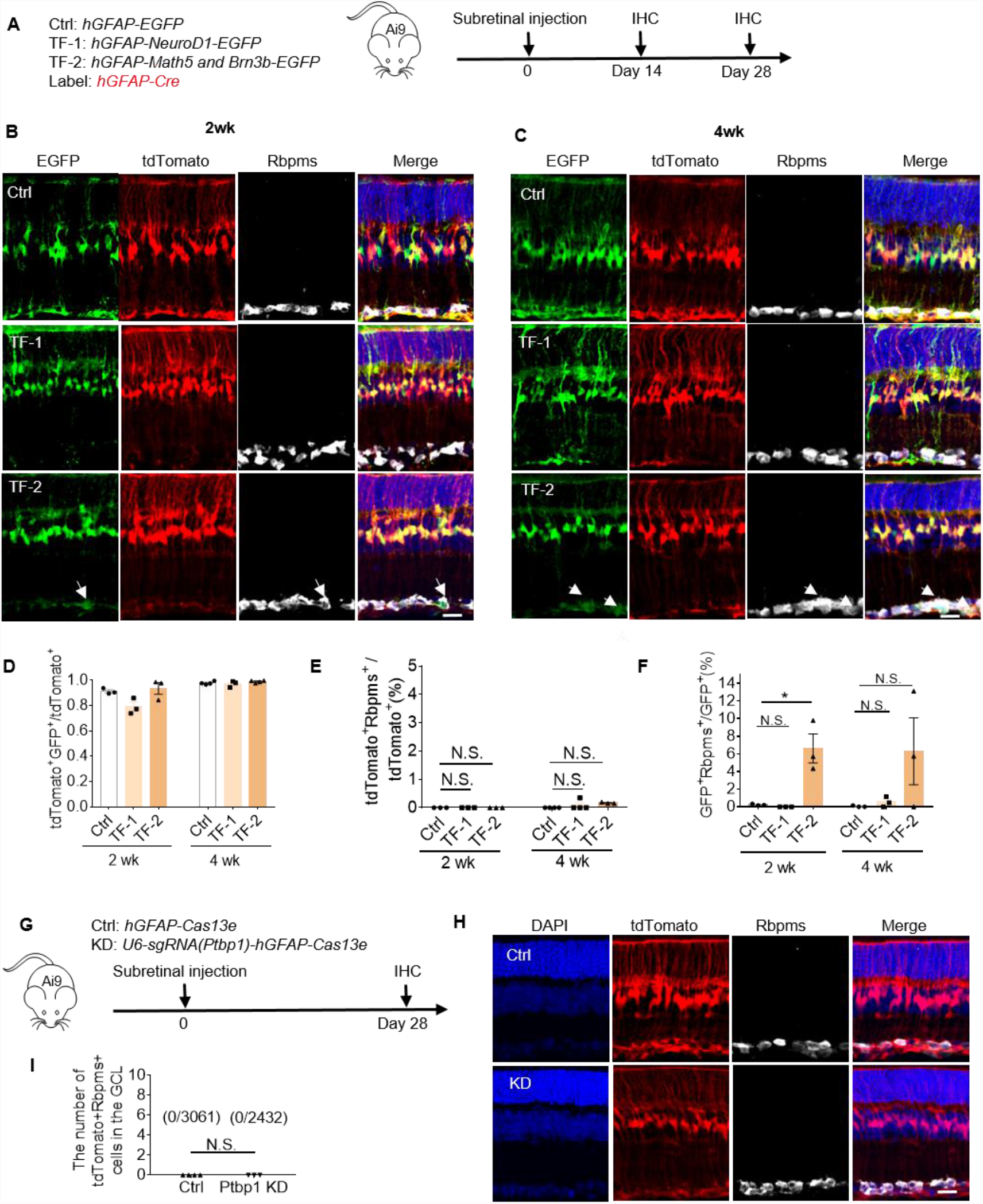
The *ΔWPRE* system fails to trace reprogrammed RGC mediated by TFs overexpression or knockdown. (A) Schematic diagram of vectors for gene transfer and injection process. (B-C) Representative images of GFP and tdTomato co-labeling with the RGC marker Rbpms at 2 weeks (B) and 4 weeks (C) after injection. Arrows show GFP^+^Rbpms^+^ cells. Scale Bar, 50 μm. (D) The ratio of tdTomato^+^GFP^+^ MG among total tdtomato^+^ cells in each eye section at weeks and 4 weeks after injection. (E) The ratio of tdTomato-labeled RGC among total tdtomato^+^ cells. (F) The ratio of GFP-labeled RGC among total GFP^+^ cells. (G) Schematic diagram of vectors for Ptbp1 knockdown and injection process. (H) Representative images of tdTomato co-labeling with the RGC marker Rbpms. Scale Bar, 50 μm. (I) The ratio of tdTomato-labeled RGC in the GCL among total tdTomato^+^ cells.

To further validated *AAV9-hGFAP-Cre-ΔWPRE* reporter system, we traced *in vivo* MG conversion to photoreceptor cells (RP) with the *ΔWPRE* system. We overexpressed Nrl, Crx and Otx2 following MG proliferation mediated by β-catenin to induce MGs into PRs, as reported in previous work (Yao et al., 2018) (Figure 5A). We traced intermediated cells, which re-entered cell cycle and performed an asymmetric cell division, and reprogrammed RP cells at 2 weeks following injection (Figure 5B), as previous work reported (Yao et al., 2018). We also detected 8.5% tdTomato^+^ RPs among total tdTomato^+^ cells (45/568) at 4 weeks. It suggested the label system can trace MG reprogramming. In addition, when we overexpressed Ascl1, we detected a large proportion of tdTomato^+^ cells in the ONL (4.3 folds in the INL) (Figure 5E-G). We noticed that GFP, used to label Ascl1 overexpression, is not expressed in most of tdTomato^+^ cells. As a result, these tdTomato^+^ cells in the ONL were not reprogrammed cells but resident cells. This suggests Ascl1 overexpression induced non-specific tracing of the *AAV9-hGFAP-Cre-ΔWPRE*.

**Figure 5.**
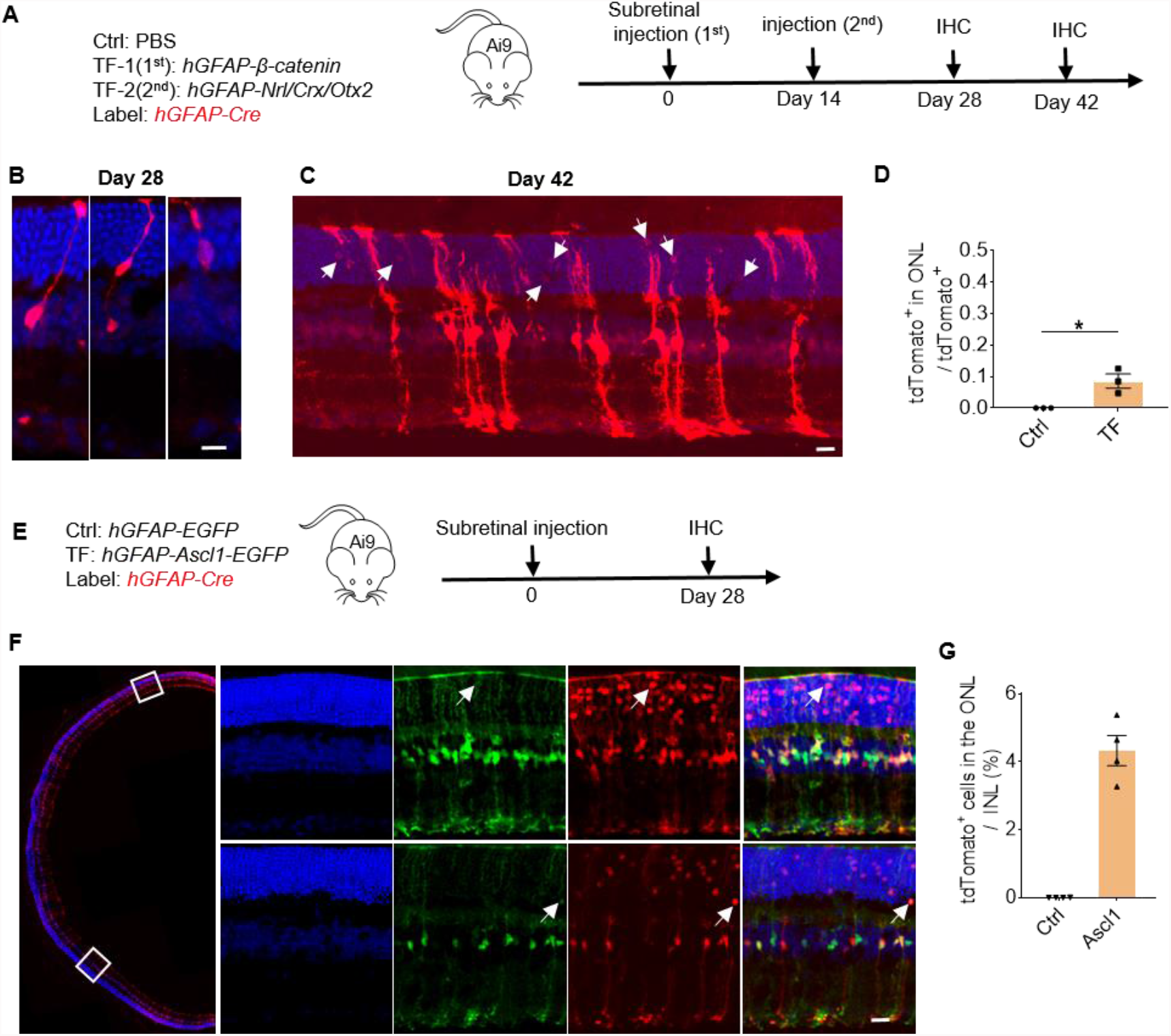
The *ΔWPRE* system labels conversion of MG to PR neuron mediated by Nrl/Crx/Otx2. (A) Schematic diagram of vectors for gene transfer and injection process. (B) Representative images of tdTomato labeled intermediated cells (left and middle) and mature phororeceptor (PR, right) neurons at 2 weeks after Nrl/Crx/Otx2 injection. (C) Representative images of tdTomato-labeled PR neurons at 4 weeks. (D) The ratio of tdTomato-labeled PR neurons among all tdTomato^+^ cells at 4 weeks. (E) Schematic diagram of Ascl1 overexpression and injection process. (F) Representative images of tdTomato-labeled and GFP-labeled PR neurons at 4 weeks. Arrows show tdTomato^+^GFP^+^ cells in the ONL. (G) The ratio of tdTomato-labeled PR neurons in the ONL to INL.

## Discussion

In this study, we developed a relatively straightforward and accessible mouse MG labeling system in which AAV9-based *hGFAP-Cre-*Δ*WPRE* was applied in Ai9 transgenic mice. This system can label MG efficiently and specifically without obvious mislabeling of other functional neurons, including RGC, bipolar and horizontal cells.

Selecting an appropriate system for tracking MG in vivo is essential for investigations of MG reprogramming. While the introduction of reporters by crossing with a transgenic Cre mouse line provides more consistent and stable labeling than AAV-based systems, the labeling efficiency may be low. In this study, we found that introduction of the *mGFAP-Cre* transgenic construct by conventional crossing failed to label the majority of MGs, unlike that reported for astrocytes in brain tissues (Garcia et al., 2004). Mouse lines with inducible Cre, such as CreER, are considered more stringent than constitutive Cre lines, but the labeling efficiency is dependent on tamoxifen dose (Hu et al., 2020), and was shown to exhibit particularly low effectiveness in tracking MG in other studies (Jorstad et al., 2017; Ueki et al., 2015). Additionally, tamoxifen administration reportedly impairs neurogenesis (C. M. Lee et al., 2020) and can adversely affect the activity and specificity of promoters driving Cre (Denk et al., 2015; Donocoff, Teteloshvili, Chung, Shoulson, & Creusot, 2020).

Considering the heterogeneity of MGs including many subtypes (Menon et al., 2019; Roesch et al., 2008), conclusions regarding the ability of MG to differentiate into other neuronal types require validation by labeling and detection of all MG subtypes. However, the labeling efficiency of MG by AAV-based reporters primarily depends on its transfection efficiency which can be affected by AAV tropisms, enhancer elements in the Cre vector (such as *WPRE*), and injection methods. The AAV1, 2, 5, 8, 9 and ShH10 serotypes with an eGFP reporter driven by CMV or CAG promoters have been shown to transfect MG using intravitreal or subretinal injection (Klimczak, Koerber, Dalkara, Flannery, & Schaffer, 2009; Lebherz, Maguire, Tang, Bennett, & Wilson, 2008; Surace et al., 2003). We compared the transfection patterns of these serotypes using an *hGFAP-Cre* vector and found that AAV1, AAV2, or AAV5 were low effective for MG transfection. Other recent work showed variable specificity by different serotypes for labeling astrocytes in the brain (Wang et al., 2021), which were in line with our results in the retina.

In addition, high specificity for tracking MG is indispensable for exclusion of false-positive results in investigation of MG conversion to neurons. Until now, the efficiency and specificity of AAV-based labeling of astrocytes remains controversial for study of neurogenesis and reprogramming, especially in the brain (Thanh Hoang et al., 2021; Wang et al., 2021). We found that all serotypes exhibited varying degrees of leakage to RGCs in the retina. We further improved the vector by removing the *WPRE* element and found that labeling efficiency was not obviously affected, but the proportion of non-specifically labeled cells decreased by at least 40-fold that of systems which include *WPRE*, substantially lower than that of other reported tracking systems for astrocytes in the brain (Srinivasan et al., 2016; Wang et al., 2021). Moreover, the system lacking *WPRE* exhibits a consistently low non-specific labeling ratio across a wide dose range, from 1×10^8^ to 1×10^10^ vg per eye, suggesting its potential reliability for tracking MG.

Using the AAV-based *hGFAP-Cre-*Δ*WPRE* system, we re-examined MG programming mediated by transcription factors reported by previous works and detected some false-positive results. First, we found overexpression of Math5/Brn3b cannot induced conversion of MG to RGC with the Δ*WPRE* system, and previous reported positive reprogrammed RGCs actually are some resident RGCs labeled by the hGFAP-GFP system (Xiao et al., 2021). This suggested that the *hGFAP-GFP* system is an improper label system for tracing conversion of MG to RGC mediated by Math5/Brn3b. However, how Math5/Brn3b mediated GFP expression in RGCs in *hGFAP-Math5/Brn3b-P2A-GFP* still be unclear and further study needs to be performed. Besides, we also failed to trace conversion of MG to RGC mediated by Ptbp1 knockdown with Cas13X.1. In previous work, the researchers used the *AAV9-hGFAP-Cre-WPRE* reporter system to trace MG lineage (Zhou et al., 2020), but we found it was not a high specific system for labeling MG in this study. As a result, other efficient and specific labeling strategies still need to be developed to double check MG programming mediated by Ptbp1 knockdown. In addition, we further verified conversion of MG to PR following Nrl/Crx/Otx2 overexpression. The intermediated and mature RPs were traced with the Δ*WPRE* system as previous work reported (Yao et al., 2018), although reprogramming efficiency is relatively low. This result suggests the Δ*WPRE* system is available to trace MG reprogramming. However, the specificity of this AAV-mediated reporter system depends on MG-specific Cre expression driven by the GFAP promoter. If the manipulated gene of interest affects GFAP promoter activity, its overexpression or knockdown in other cell types, such as neurons, may activate the GFAP promoter and initiate Cre expression in those neurons, in turn affecting its specificity for MG. In this study, Ascl1 may regulate the *hGFAP* promoter and result in leakage of tdTomato. As previous work reported, NEUROD1 may function as a *cis*-regulator of the *hGFAP* promoter (Wang et al., 2021). Under this condition, other tracking strategies, such as labeling neurons with retrograde virus (Wang et al., 2021), should be combined with this method to further validate the results. In all, our AAV9 *hGFAP-Cre-*Δ*WPRE* system could offer conveniences for fundamental research in both higher and lower vertebrate eye development, as well as the potential for development of MG-targeted therapeutics. At the same time, more cautions should be taken when we use *in vivo* MG lineage-tracing to detect reprogramming.

## Acknowledgements

We thank Dr. Jiawei Zhou and Dr. Zilong Qiu for offering transgenic *mGFAP-Cre* and *Aldh1l1-Cre* mouse line, Xinde Hu from Haibo Zhou laboratory for suggestions on plasmid construction, and the Optical Imaging facility, Y. Wang, Y. Zhang and Q. Hu in ION. We also thank Isaac V. Greenhut for discussions and comments on this manuscript. This work was supported by the Basic Frontier Scientific Research Program of Chinese Academy of Sciences from 0 to 1 original innovation project (ZDBS-LY-SM001), the R&D Program of China (2017YFC1001300 and 2018YFC2000100), the CAS Strategic Priority Research Program (XDB32060000), the National Natural Science Foundation of China (31871502, 31925016, 91957122, 31901047, 82001355), the Shanghai Municipal Science and Technology Major Project (2018SHZDZX05), the Shanghai City Committee of Science and Technology Project (18411953700, 18JC1410100, 19XD1424400 and 19YF1455100) and the International Partnership Program of Chinese Academy of Sciences (153D31KYSB20170059).

## Author contributions

Y.G., K.F. and H.Y. conceived the project. Y.G., K.F., Z.Y. and H.Z. designed and conducted experiments. W. W. and G.G. assisted with virus package. D.X. and H.Z. performed cryostat section for retinas. N.Z., Q.W., M.C and E.Z. assisted with animal experiments. H.Y. designed experiments and supervised the whole project. Y.G., K.F., Z.Y. and H.Y. wrote the paper.

## Figures and Figure legend

**Figure S1.**
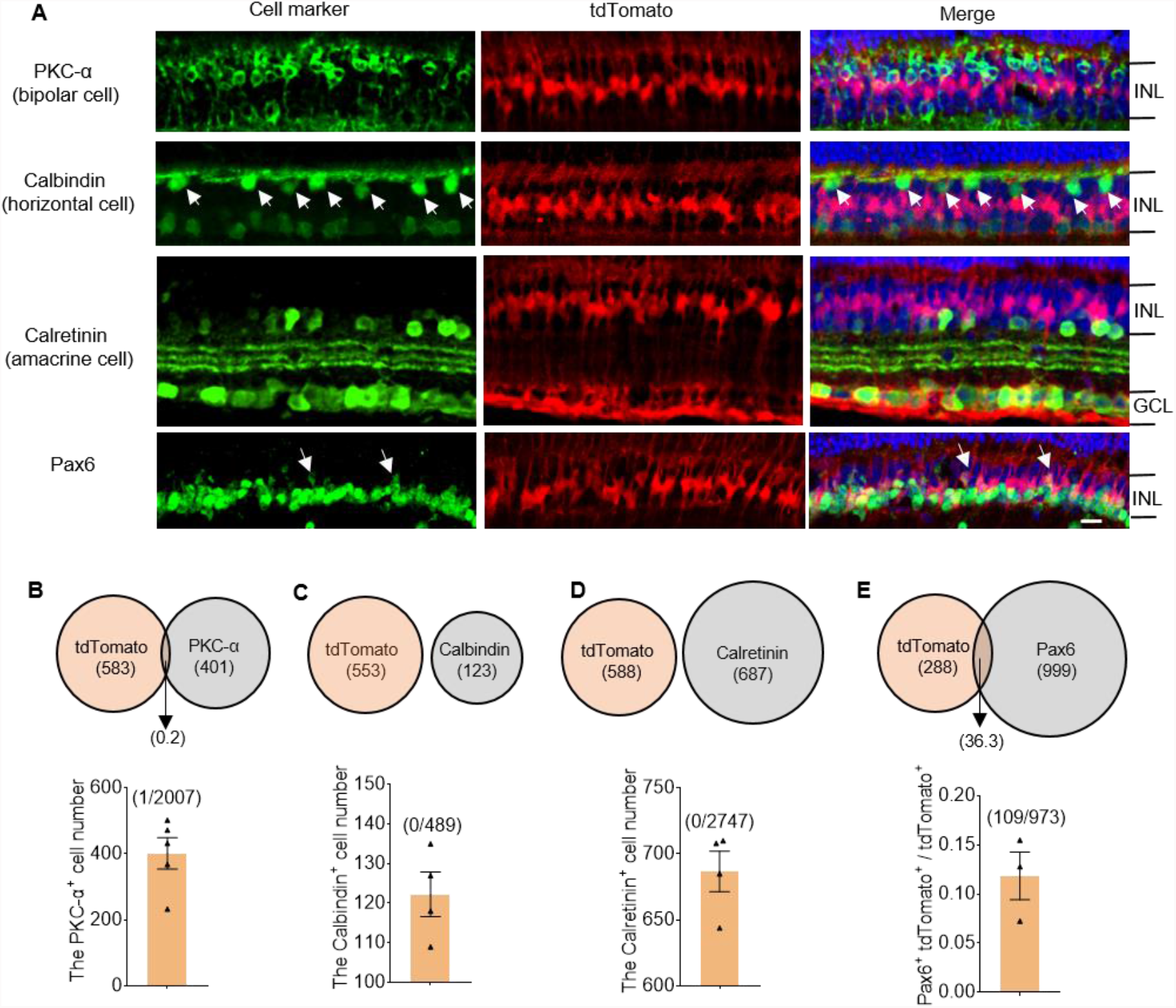
Reduced leakage in the *ΔWPRE* system related to Figure 3. (A) Co-localization of tdTomato and bipolar cell marker PKC-α, horizontal cell marker Calbindin and amacrine cell marker Calretinin and Pax6. Arrows show horizontal cells. INL, inner nuclear layer. Scale Bar, 50 μm. (B-E) Venn diagram of average tdTomato reporter and cell markers in each section (up); the number of PKC-α^+^ (C), Calbindin^+^ (C), Calretinin ^+^ (D) and the ratio of tdTomaot^+^Pax6^+^ to tdTomato^+^(E) cells in each eye section infected at 10^9^ vg/eye (down). PKC-α marked bipolar cells. Calbindin marked horizontal cells, amacrine cells and RGCs in mice but could distinguish horizontal cells, pointed out by arrows. Calretinin and Pax6 mainly marked mouse amacrine cells. Brackets in panel (B-D) show the number of both tdTomato and cell marker positive cells among total cell marker positive cells. Brackets in panel (E) show the total number of tdTomato^+^Pax6^+^ and tdTomato^+^ cells. n=4∼5 retinas for panel B-E. All values are presented as means ± S.E.M.

